# Supervised Factorization to Associate Spatial Transcriptomics with Complementary Molecular Readouts

**DOI:** 10.1101/2025.09.28.679034

**Authors:** Faria Binta Awal, Robia G. Pautler, Md. Abul Hassan Samee, M Saifur Rahman

**Affiliations:** Department of Computer Science and Engineering, Bangladesh University of Engineering and Technology (BUET), Dhaka, 1000, Bangladesh; Department of Integrative Physiology, Baylor College of Medicine, Houston, Texas, United States

## Abstract

Spatial Transcriptomics enables studying gene expression data within spatial context of tissues. Yet understanding how spatial molecular phenomena influence transcriptional patterns remains a key challenge. We propose a novel supervised Non-negative Matrix Factorization (NMF) framework, where supervision is selectively and explicitly applied to guide the learning of a supervised spatial factor. This distinguishes our method from prior approaches by enforcing spatial alignment only on a targeted component of the factorization, enabling biologically interpretable associations between gene expression and spatial molecular events. This approach also enables the identification of genes whose expression patterns are spatially correlated with molecular events of interest. Applied to datasets involving Alzheimer’s Disease (AD) and Myocardial Infarction (MI), our method successfully discovered supervised spatial factor associated with disease related signal. In the case of Alzheimer’s Disease (AD), we have presented a spatial decay model to represent how the influence of amyloid-beta plaque signals diminishes with distance, and used this as a supervision signal during matrix factorization. Applied across both disease contexts, our method successfully identified biologically meaningful gene sets associated with disease progression. By ranking genes based on their contribution to the supervised spatial factor, the framework highlights candidate genes potentially involved in disease-related processes.

## Introduction

Spatial Transcriptomics (ST) has emerged as a transformative technology for understanding tissue biology by preserving the spatial context of gene expression within native tissue architecture. By analyzing transcriptomic data in its spatial context, ST enables the discovery of gene expression modules: coordinated sets of genes that define distinct biological processes and cellular states. These spatially resolved modules provide unprecedented insights into disease mechanisms, particularly in understanding how cells respond to pathological signals and the molecular changes that contribute to disease progression.

A critical next frontier in spatial biology is the joint analysis of ST data with complementary spatial modalities collected from the same or adjacent tissue sections. Such multimodal approaches—combining ST with immunofluorescence, histopathology images, or disease-specific markers like amyloid plaques—promise to reveal how gene expression modules relate to observable phenotypes and pathological features. For instance, in Alzheimer’s disease, the hallmark neuropathological features [15, 3, 2, 22] include widespread accumulation of extracellular A*β* (amyloid beta) plaques. So, aligning ST data with A*β* plaque imaging could identify gene modules that respond to or drive plaque formation, while in cardiac diseases, tissue damage markers could guide the discovery of injury-associated expression programs. However, despite the clear potential of these multimodal analyses, we currently lack computational methods that can effectively model ST data while incorporating supervision from aligned imaging modalities.

Current spatial transcriptomics platforms, such as, the GeoMx Digital Spatial Profiler by NanoString, enable researchers to profile gene expression within manually defined regions of interest (ROIs), allowing for comparative analysis between spatially distinct zones (e.g., plaque-proximal vs. plaque-distant regions) [14]. However, these approaches rely on user-defined ROI selection and descriptive comparisons, lacking computational methods that can systematically integrate continuous supervision signals from imaging modalities to automatically discover gene expression modules associated with pathological features.

Existing approaches for ST analysis, such as Non-negative Matrix Factorization (NMF) [20] and its spatial extension Non-negative Spatial Factorization (NSF) [29], operate in an unsupervised manner. When applied to a non-negative data matrix *X*, where rows correspond to observations and columns correspond to features, NMF breaks it down into two non-negative matrices: *W*, often referred to as the basis or factor matrix, and *H*, known as the coefficient matrix (Figure 1). This factorization helps us to understand the complex dataset by uncovering the latent factors.

**Fig. 1.**
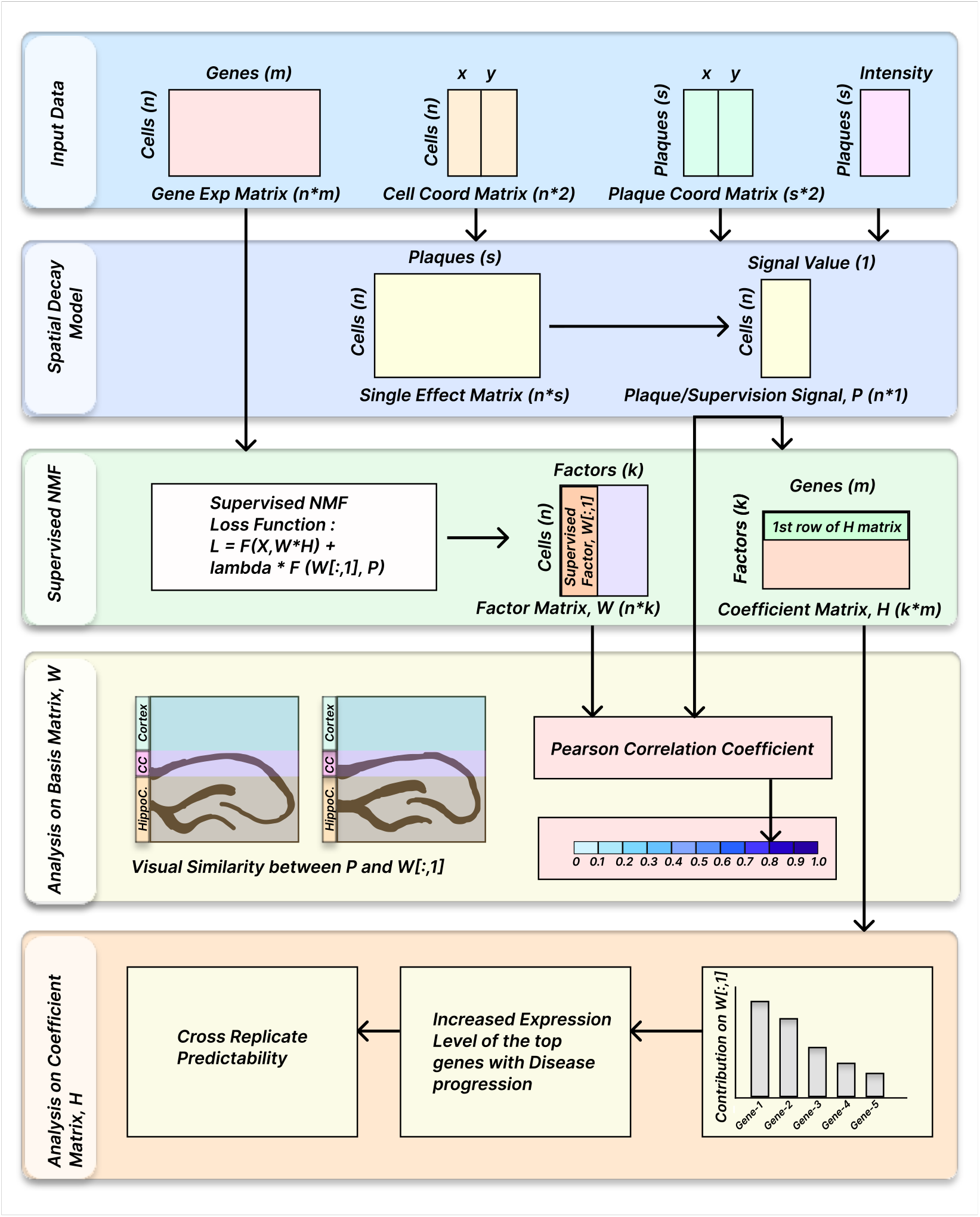
Overview of the SuNSTAMP Pipeline, illustrated for the AD dataset. The same pipeline was also applied to the MI dataset except the spatial decay model stage. The pipeline consists of five key stages: (1) Preprocessing input data, including gene expression matrix, cell coordinates, and data from a second modality (A*β* plaques) (2) Spatial Decay Model, which computes a spatially-aware supervision signal by modeling the influence of A*β* plaques on cells (3) Supervised NMF, where gene expression matrix is factorized with supervision signal applied to a single latent factor; (4) Analysis of *W* matrix, evaluating the alignment between the supervised factor and the A*β* supervision signal and (5) Analysis of *H* matrix, interpreting gene contributions towards the supervised factor and performing cross-replicate validation for biological insights and generalizability.

While these methods successfully decompose expression data into interpretable factors that correspond to spatial domains, they cannot leverage external signals from the complementary modalities to guide factorization. This limitation prevents researchers from directly identifying gene modules that are specifically associated with pathological features or other biologically relevant signals captured in companion imaging data.

Here, we address this methodological gap by introducing SuNSTAMP (**Su**pervised **N**MF to define **S**patial **T**ranscriptomics Modules underlying **A**dditional **M**olecular **P**henotypes), a novel framework that extends NMF to incorporate supervision from aligned spatial modalities. Our key innovation is the targeted application of supervision to a single factor within the NMF decomposition (Supplementary Figure S1), enabling this factor to capture gene expression patterns specifically associated with the external signal while preserving the unsupervised discovery of other biologically relevant factors. Through this design, SuNSTAMP not only identifies genes whose spatial expression correlates with pathological features but also provides a quantitative ranking of their contribution to the supervised phenotype.

We demonstrate SuNSTAMP’s utility across two distinct disease contexts with different pathological characteristics - Alzheimer’s Disease (AD) and Myocardial Infarction (MI). AD reflects neurodegenerative pathology with spatially heterogeneous plaque deposition, while MI involves acute tissue damage and localized inflammatory response. In Alzheimer’s disease, we use A*β* plaque intensity as supervision to discover gene modules associated with plaque proximity. Cross-replicate validation using the learned gene weights from factorization and a compact gene set demonstrates exceptional predictive capability, achieving (AUC-ROC ≥ 0.90) for predicting plaque locations in independent biological replicates—a predictive capability not commonly found in traditional matrix factorization methods. In myocardial infarction, we leverage SPaSE-derived [26] tissue damage scores to identify expression programs linked to cardiac injury (Pearson r = 0.817). These applications showcase how SuNSTAMP provides a general framework for multimodal spatial analysis, enabling researchers to systematically discover gene expression modules that underlie observable tissue phenotypes and advance our understanding of spatially resolved disease mechanisms.

## Results

### Overview of SuNSTAMP

Our proposed framework (Figure 1) utilizes a supervised Non-negative Matrix Factorization (NMF) approach for the quantitative analysis of the contribution of features towards a targeted supervised latent factor. The input to the model is a two-dimensional data matrix *X*_*n×m*_, where *n* represents the number of observations and *m* denotes the number of features.

In our setting, *X* denotes gene expression matrix where each observation corresponds to a spatial unit — such as a spot, tissue domain, or individual cell and each feature corresponds to the expression level of a specific gene. Thus, each row of *X* captures the transcriptional profile of a specific spatial location. In addition to gene expression, SuNSTAMP integrates spatial coordinates for each observation and incorporates an additional spatial modality—such as pathological marker intensity (e.g., A*β* plaques for Alzheimer’s disease or infarct severity in Myocardial Infarction)—characterized by location-specific values. This second modality serves as a biologically meaningful supervision signal to guide the decomposition process.

The model factorizes *X*_*n×m*_ into two non-negative matrices:

- *W*_*n×k*_: A basis or factor matrix, where *k* represents the number of latent factors.
- *H*_*k×m*_: A coefficient matrix, capturing the contributions of the features (genes) to the latent factors.

The supervision is applied to the first column of *W*, aligning it with the targeted supervision signal, thereby making it a targeted supervised factor (Supplementary Figure S2 a,b). In general, supervision applied to a specific latent factor results into the modifications to the corresponding row of matrix *H* (Supplementary Figure S1).

The factorization process minimizes a custom loss function that combines the reconstruction error of the matrix *X* with a supervision alignment. The mathematical formulation of the loss function is as follows:

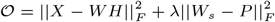

where *W*_*s*_ is the supervised factor of *W, P* is the supervision signal, and *λ*, a regularization parameter, controls the trade-off between reconstruction accuracy and adherence to the supervision signal.

To select the minimal sufficient number of latent factors and best-fit regularization parameter *λ*, we use a k-fold (k = 10) nested cross-validation strategy. The minimal sufficient rank is determined based on diminishing reconstruction improvements, and validated using the Wilcoxon rank-sum test [30] to confirm that increasing *k* beyond a particular point yields no significant gain.

To assess how well the supervised factor (first column of *W*) captures the target signal, we compute the Pearson Correlation Coefficient (PCC) [24] between *W*_*s*_ and *P*. For qualitative insight, we also visualize both the supervision signal and the supervised factor across spatial coordinates.

Finally, the first row of *H* identifies the features (genes) most associated with the supervised factor. We consider the genes with the highest weights as top contributors or top ranked genes.

Biological datasets often include multiple replicates at the same time point, raising the question of how well the learned factors generalize across replicates. To assess this, we evaluated the predictive capacity of plaque proximity of the top-ranked genes—identified from one replicate—on a different replicate within the same time point in AD. Specifically, we computed a weighted predictive score for each cell in the target replicate, using the weights of the top genes from the source replicate. We measured how well the predictive score corresponded to the spatial distribution of disease-relevant pathology using AUC analysis with plaque proximity as the target. This cross-replicate evaluation demonstrates that SuNSTAMP captures robust and transferable biological insights, providing strong evidence of generalizability across biological replicates within same time point.

Together, SuNSTAMP enables a targeted, spatially-informed decomposition of gene expression data guided by an additional disease-relevant modality. By focusing supervision on a single latent factor, the framework offers clarity, interpretability, and adaptability across diverse biological contexts.

One of the challenging applications of SuNSTAMP is to compute supervision or pathology-derived signals in a principled way, particularly when dealing with spatially heterogeneous deposition patterns that exhibit neighborhood proximity effects. To address this challenge, we propose a Spatial Decay Model (SDM) (Section 2.2) to generate such signals when pathology intensity gradients are spatially relevant, such as the characteristic spreading patterns observed in Alzheimer’s disease pathology.

### Spatial Decay Model (SDM)

Many biological systems exhibit spatially heterogeneous patterns where localized pathological features influence neighboring regions with varying intensity based on proximity. In such contexts, the biological impact of these features extends beyond their immediate location, creating gradients of influence that decay with distance. Traditional approaches often treat spatial units as isolated entities, potentially overlooking the broader spatial context and cross-regional influences that are crucial for understanding disease progression and cellular states.

We introduce the Spatial Decay Model (SDM), a quantitative framework for modeling how localized pathological features influence cellular states across spatial domains. The model incorporates two key principles: (1) feature intensity weighting, where larger or more intense pathological features exert greater influence, and (2) spatial decay functions that model how this influence diminishes with increasing distance from the source.

For a cell *C* with coordinates (*c*_*x*_, *c*_*y*_) and a pathological feature *F* with coordinates (*f*_*x*_, *f*_*y*_) and intensity measure *I*_*f*_, the impact value of feature *F* on cell *C* is calculated as:

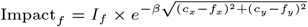

where *β* is a scaling parameter that controls the rate of spatial decay, and *I*_*f*_ represents the intensity or size of the pathological feature.

Since biological systems often contain multiple pathological features that can collectively influence cellular states, we aggregate the individual impact values across all features (Figure 2a). Following the principle that complex disease manifestations often result from additive effects of multiple contributing factors [31], the total impact score for a cell is computed as:

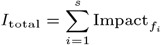

where *s* is the number of pathological features in the dataset. This summation provides a supervision signal for each cell, forming a supervision matrix *P* with dimensions *n ×* 1 where *n* is the total number of cells.

**Fig. 2.**
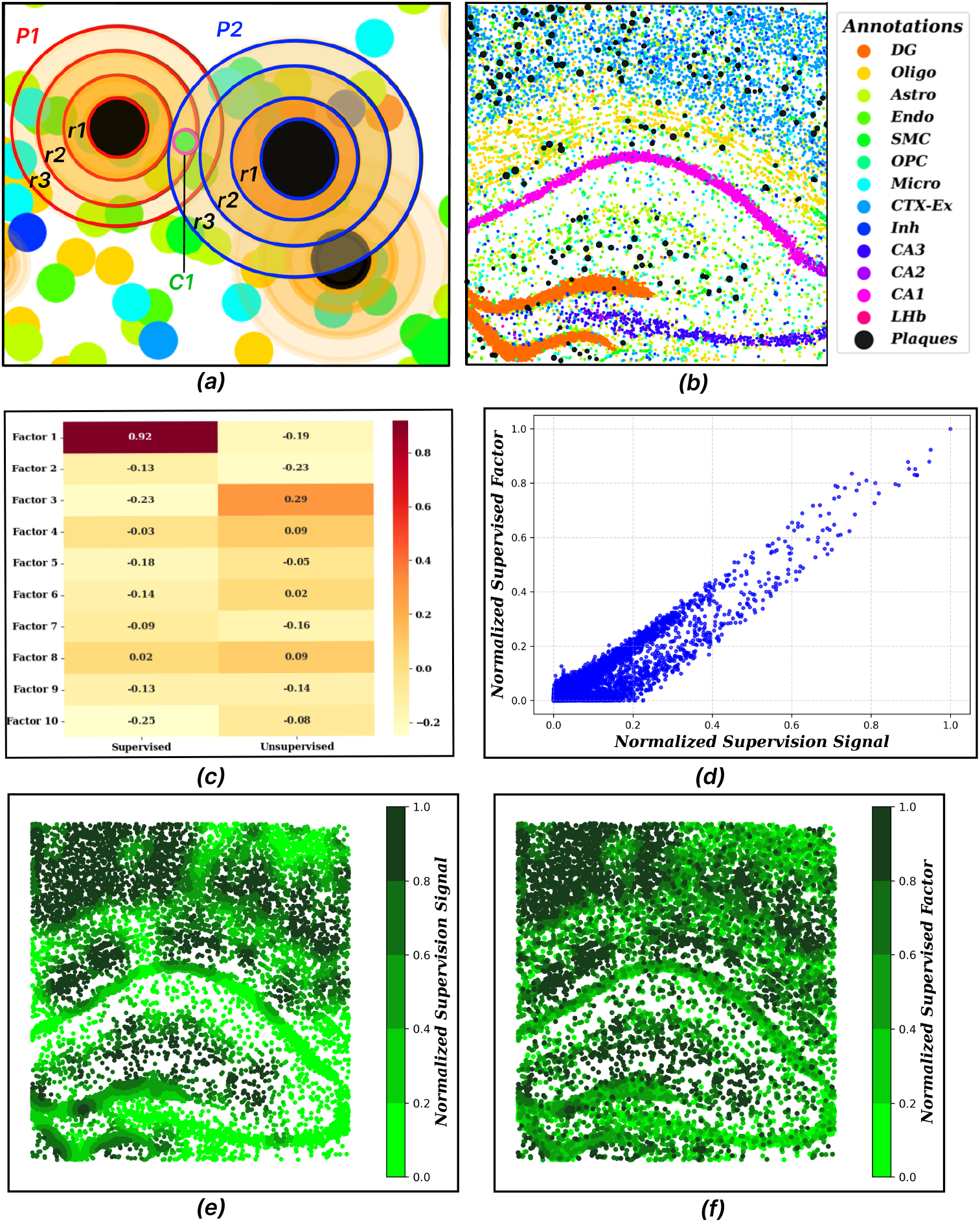
*(a)* This zoomed-in image illustrates the decay effects from three plaques, with concentric circles representing different regions of influence. These regions are labeled as *r*_1_, *r*_2_, and *r*_3_, where *r*_1_ has the highest effect, *r*_2_ has a moderate effect, and *r*_3_ has the lowest effect. The cell *C*_1_ lies in the *r*_2_ region of plaque *P*_1_ and the *r*_3_ region of plaque *P*_2_, highlighting the need for aggregation across multiple plaques. *(b)* Spatial distribution of 13 cell types and A*β* plaques in the 13-month-old diseased Alzheimer’s disease (AD) mouse brain. Each cell type is represented with a distinct color, showing their spatial organization across the brain region with A*β* plaque locations and intensities overlaid. *(c)* Pearson correlation coefficients between each of the 10 factors and the supervision signal. The first column shows results from SuNSTAMP, where the first factor (supervised factor) achieves a correlation of 0.92. The second column shows results from standard NMF (unsupervised), with the best factor reaching only to 0.29. *(d)* Scatter plot showing the relationship between normalized supervision signal (derived from spatial decay model of A*β* plaques) and normalized supervised factor (learned through SuNSTAMP). Points are concentrated near the origin and follow an approximately 45-degree trajectory. *(e)* Spatial distribution of the supervision signal derived from the Spatial Decay Model, showing theoretical pathological influence based on A*β* plaque proximity and intensity. *(f)* Spatial distribution of the normalized supervised factor learned by SuNSTAMP from gene expression data. Both visualizations use identical 5-bin quantile-based color mapping (bright green to dark green) to enable direct comparison. The remarkable spatial concordance between theoretical pathological influence and learned factor values demonstrates that SuNSTAMP successfully identified cells in pathologically relevant anatomical regions.

#### Application to Alzheimer’s Disease

To demonstrate the utility of SDM, we apply it to model the spatial influence of amyloid-beta (A*β*) plaques in Alzheimer’s disease. Several studies show that A*β* plaques disrupt nearby cells in AD-affected brains, accelerating disease progression in their vicinity [28]. Intuitively, cells situated closer to larger plaques should indicate more advanced disease states, as proximity to plaques accelerates disease progression.

Previous approaches, such as the work by Chen et al. [7], used hierarchical clustering alongside Weighted Gene Co-expression Network Analysis (WGCNA) to identify Plaque-Induced Gene networks (PIGs). However, their spot-centric approach treats each tissue spot as an isolated spatial unit, restricting the influence of A*β* plaques to the specific spot where they are detected (Supplementary Figure S2 c). This rigid approach overlooks the fact that genes associated with plaque pathology may still be expressed in spots with low or no detectable plaque, influenced by nearby regions with higher plaque intensity.

In our AD application, we model plaque intensity *I*_*f*_ based on the estimated volume of the plaque, approximating it as a sphere:

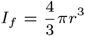

where *r* is the plaque radius. This approach enables us to spatially smooth the A*β* plaque intensity across neighboring spots, more accurately capturing the biological influence of plaques on surrounding tissue (Supplementary Figure S2 d).

### SuNSTAMP discovers a Plaque Associated Supervised Factor in AD dataset

In this section, we demonstrate SuNSTAMP using the high-resolution spatial transcriptomics dataset from the STARmap PLUS [32] platform, obtained from an Alzheimer’s disease (AD) mouse model. The analysis for the myocardial infarction (MI) dataset is presented in a later section. The AD dataset analyzes gene expression alterations at different disease stages, specifically in 8-month-old and 13-month-old mice. The dataset comprises 9192 transcriptomic profiles w ith 2 766 relevant genes from AD-related databases. Each profile includes spatial coordinates, cell types, and gene expression levels. A*β* plaque details - such as the coordinates and radius - are provided separately in the same coordinate system, allowing us to explore the relationship between gene expression changes and plaque load measurements (Figure 2b). Eight rounds of in situ sequencing and one round of post-sequencing imaging were performed to locate AD-related pathology, particularly Amyloid beta (A*β*) plaques in coronal sections of the brains of TauPS2APP mice and control mice. In total, 13 distinct cell types are distributed across the three major brain regions—Cortex, Corpus Callosum (CC), and Hippocampus. The full list of cell type acronyms, their corresponding names, and distribution counts for a 13 month old AD mouse model are provided in Supplementary Table S1.

To demonstrate the effectiveness of our supervised factorization approach, we computed the Pearson Correlation Coefficient (PCC) [24] between the learned supervised factor and the supervision signal derived from the spatial decay model of A*β* plaque signals (P). The resulting correlation is 0.92 (Figure 2c), indicating a strong linear relationship between the supervised factor and the target pathological signal. This significantly o utperforms t he b est correlation obtained by any single factor in the unsupervised factorization, which was only 0.29, demonstrating the clear advantage of incorporating spatial pathology information into the factorization process. The scatter plot visualization (Figure 2d) reveals a concentrated distribution of points near the origin with a clear upward trend following approximately a 45-degree trajectory, confirming the strong positive correlation. This high correlation ensures that SuNSTAMP successfully guided the factorization process by aligning the supervised factor with the supervision signal, whereas unsupervised NMF fails to produce any factor that correlates meaningfully with the target pathological signal. The guided factorization process thus enhances the interpretability of the supervised factor in the basis matrix *W*, directly linking it to spatial and pathological attributes relevant to Alzheimer’s disease progression.

To complement the quantitative evaluation, we visualize the spatial distribution of the supervision signal (Figure 2e) alongside the supervised factor (Figure 2f) in the 13-month-old mouse brain, primarily for interpretability. The supervised factor map reveals remarkably similar spatial patterns, with high factor values concentrated in the same anatomical regions where plaque pathology is most pronounced.

### SuNSTAMP Recapitulates Known Biology and Uncovers New Insights

After confirming that the supervised factor aligns well with the supervision signal, we analyze the contribution of individual genes to this factor. Since the supervision signal was designed to reflect disease-associated features—specifically, A*β* load measurements — the first row of the *H* matrix reveals how strongly each gene contributes to this factor. By sorting these contribution values in decreasing order, we obtain a ranked list of genes, highlighting their relevance to the supervised factor and, by extension, their association with plaque load.

The top 10 genes from the ranked gene list consists of *C1qa, Ctsb, Cst3, Tmsb4x, Apoe, Hexb, Gfap, Cd9, Cst7, Gna14* in decreasing order of their contribution value towards the supervised factor. In Figure 3a, the numeric values above each bar represent the genes’ exact contributions or weights. For instance, *C1qa* has a contribution value of 0.27, while *Gfap* has a value of 0.15, indicating that *C1qa* contributes approximately 1.8 times more than *Gfap* towards the supervised factor and the associated plaque effects.

**Fig. 3.**
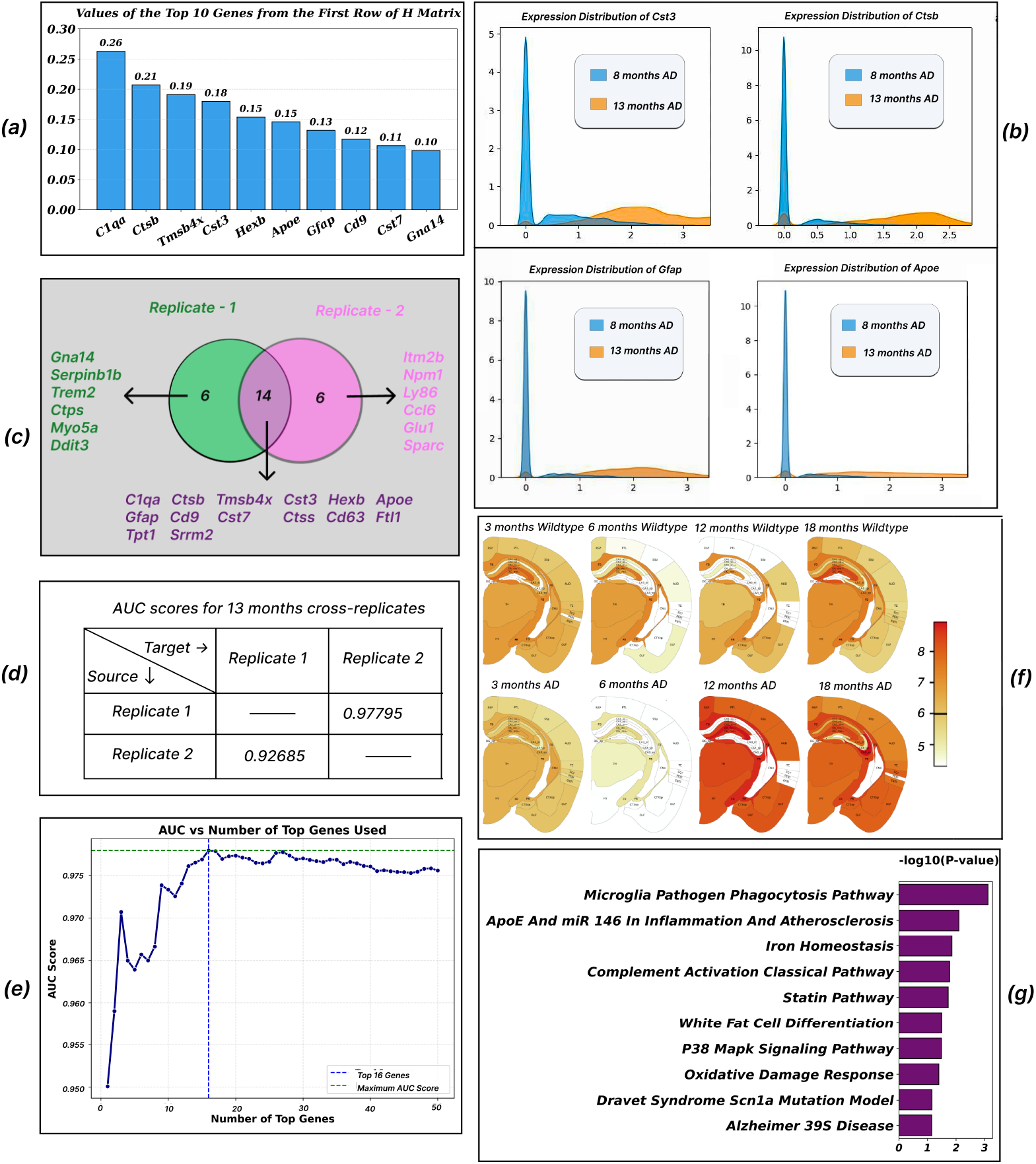
*(a)* Top 10 genes contributing the most to the supervised factor found in 13 months old AD mouse brain (replicate-1), sorted in decreasing order of their values in the first row of the *H* matrix. *(b)* Expression distribution of four plaque-associated top genes (*Cst3, Ctsb, Gfap, Apoe*) in 8- and 13-month-old AD mouse models, visualized using Kernel Density Estimation (KDE). The x-axis represents gene expression level (log-normalized counts), and the y-axis represents probability density, where the area under each curve equals 1 for each age group. For all four genes, the 13-month distribution curves show a noticeable rightward shift relative to the 8-month curves, indicating that a larger proportion of cells exhibit higher expression levels at the later disease stage. This pattern suggests progressive gene upregulation with increasing A*β* plaque burden. *(c)* Venn diagram showing the overlap between the top 20 genes identified from two independent replicates (Replicate 1 and Replicate 2) of 13-month-old AD mouse brains. The central region shows the 14 common genes, while the outer sections list genes unique to each replicate. *(d)* Cross-replicate validation using AUC-ROC scores. Each entry shows the predictive performance when the model is trained on the source replicate (row) and tested on the target replicate (column). *(e)* AUC-ROC scores for increasing numbers of top-ranked genes (1–50). The model reaches peak performance while using top 16 genes. *(f)* Expression profile of one of the top-ranked genes - *Apoe*, across control and Alzheimer’s disease (AD) mouse brains at 3, 6, 12, and 18 months from an independent dataset [7]. The gene’s expression is visualized with control (top row) and AD (bottom row) across these 4 different time points. Warmer color (red) indicate higher expression levels. Notably, elevated expression in AD mice at 12 and 18 months supports the relevance of SuNSTAMP-identified genes in later stages of disease progression. *(g)* Statistical significance (-log10 P-value) of enriched pathways from the top 20 genes identified by the SuNSTAMP supervised factor. Key AD-related pathways including microglia phagocytosis, *ApoE* inflammation, and complement activation show significant enrichment, validating the biological relevance of the supervised factorization approach.

Several studies suggest that *C1qa* has a detrimental role in Alzheimer’s disease (AD) by promoting synapse loss[10]. The findings indicate that *C1q*-dependent mechanisms drive both astrocytic and microglial engulfment of synapses, leading to pathological synapse elimination in TauP301S mice. In the context of Alzheimer’s disease, *C1qa, Grn*, and *Ctsb* are upregulated in microglia in response to SPP1 signaling, promoting phagocytosis of synapses [9]. Increased *Gfap* levels, particularly in plasma, may reflect early astrocyte activation, potentially preceding amyloid plaque formation and influencing the progression of AD pathology [25]. Together, these observations reinforce the biological plausibility of our ranked gene list and demonstrate that the supervised factor effectively captures meaningful disease-associated signal.

We selected several key genes *Cst3, Ctsb, Gfap* and *Apoe* from the top 10 genes’ list to investigate their expression patterns across 8-month and 13-month AD mouse models using Kernel Density Estimation (KDE) [8]. Distribution plots of these genes revealed a noticeable rightward shift in their expression levels at 13 months compared to 8 months, indicating significant upregulation in response to increasing A*β* loads (Figure 3b). This upregulation re-emphasizes the potential role of these genes in the progression of Alzheimer’s disease and highlights their relevance as key contributors to plaque-associated pathology.

To assess the robustness of SuNSTAMP’s gene discoveries, we compared the top 20 genes contributing to the supervised factor in two independent replicates — both from 13-month-old mouse brain samples. Figure 3c illustrates the overlap between the gene lists using a Venn diagram.

Out of the 20 top-ranked genes from each replicate, 14 were found to be common. These shared genes are displayed in the overlapping region of the diagram, while replicate-specific genes appear in the non-overlapping areas. The presence of such substantial overlap indicates that SuNSTAMP consistently identifies biologically relevant genes across replicates, despite differences in sample variation or factorization noise.

Importantly, most of these 14 common genes—such as *C1qa, Ctsb, Tmsb4x, Cst3, Hexb, Apoe, Gfap, Cd9, Cd63* and *Cst7* —have been extensively studied in the context of Alzheimer’s disease, underscoring the biological plausibility of SuNSTAMP’s results. At the same time, the presence of less-characterized genes within this consistent set highlights promising candidates for further investigation, potentially shedding light on underexplored molecular contributors to AD pathology.

To further validate the stability of SUNSTAMP’s gene prioritization, we assessed whether the relative orderings among the 14 genes shared between replicate 1 and replicate 2 were preserved. Using pairwise comparisons, we found that 70% of all possible gene pairs maintained consistent ordering across two independent replicates in 13-months-old AD mice (Supplementary Figure S3).

This result suggests that not only does SuNSTAMP identify overlapping sets of biologically relevant genes, but it also maintains stable internal ranking among them—despite being applied to independent replicates within the same time point.

### Predictive Validation of Top AD Genes Across Replicates

To assess the predictive power of the genes identified by our model in distinguishing spatial spots near versus far from A*β* plaques, we used the learned gene weights and computed a weighted gene expression score for each spatial spot in an independent replicate, serving as a predictive score for plaque proximity. We evaluated this predictive score using AUC-ROC, treating the binary classification of each spot as either *near* or *far* from plaques. Plaque proximity was defined using a distance threshold of 30 units from the plaque boundary, with spots within this range labeled as *near* and others as *far*. To assess generalizability, we conducted cross-replicate validation: the model was trained on replicate 1 (13 months) and evaluated on replicate 2, and vice versa. In both cases, we obtained AUC-ROC scores exceeding 0.90, highlighting the reproducibility of the predictive genes across replicates. These results are summarized in Figure 3d.

We further examined how the number of top-ranked genes influences predictive performance. For both replicates, we incrementally included the top k genes (from 1 to 50) and computed the AUC-ROC at each step. As illustrated in Figure 3e, performance increases with more genes but plateaus appear after approximately 15–20 genes, suggesting a compact and robust gene set carries most of the predictive power.

These findings suggest that the top genes selected and ranked by our model are not only biologically meaningful but also reliably predictive of spatial plaque proximity across independent biological replicates.

To further validate the significance of our top-ranked genes from SuNSTAMP, we evaluated their expression patterns in an independent dataset [7] featuring gene expression profiles from wild-type (control) and Alzheimer’s disease (AD) mouse brains. This dataset spans four time points: 3, 6, 12, and 18 months, providing a comprehensive view of gene expression changes across different stages of disease progression.

Our findings reveal that the expression levels of these genes are significantly higher in AD mouse brains at 12 and 18 months compared to wild-type mice (Figure 3f). The gene expression distribution plots shown in Supplementary Figure S4 were obtained from [AlzMAP](https://alzmap.org/), an online resource provided alongside the dataset. This observation highlights the involvement of these genes in later stages of disease progression. These results reinforce the biological relevance of the top genes identified by our supervised approach and their significance in Alzheimer’s disease mechanisms.

To assess the biological relevance of the supervised factor, we also performed pathway enrichment analysis using Enrichr [18] on the top 20 highest-weighted genes identified by SuNSTAMP. The analysis revealed significant enrichment of pathways directly associated with Alzheimer’s disease pathophysiology (Figure 3g). The most significantly enriched pathway was microglia pathogen phagocytosis (P = 0.0007), followed by ApoE-mediated inflammation, iron homeostasis, and complement activation—all key mechanisms implicated in AD progression. The enrichment of microglia pathogen phagocytosis pathway in our supervised factor aligns with previous discoveries of specialized microglial clearance mechanisms in Alzheimer’s disease. Recent work has identified LC3-associated endocytosis (LANDO) as a critical process by which microglia clear A*β* aggregates, with deficient LANDO leading to increased neuroinflammation and accelerated neurodegeneration [16].

These results demonstrate that the supervised factor successfully captures genes involved in established AD-related biological processes, providing strong evidence for the biological interpretability and disease relevance of our approach.

### SuNSTAMP Signals Apply Broadly across Pathological Landscapes

To evaluate the versatility of SuNSTAMP beyond neurodegenerative contexts, we applied our framework to a spatial transcriptomics dataset of mouse hearts affected by Myocardial Infarction (MI), originally published by Calcagno et al. [4]. This dataset captures gene expression from key anatomical regions of mouse heart at multiple post-infarction time points—specifically 3 days and 7 days after MI induction, including: Remote Zone (RZ; healthy tissue), Border Zone 1 (BZ1; adjacent and stressed tissue), Border Zone 2 (BZ2; more affected tissue), and the Infarct Zone (IZ; severely damaged and fibrotic tissue). We used pathological scores derived from the SPaSE tool [26] as our supervision signal (Figure 4a), reflecting the severity of tissue damage at each spatial spot. Higher scores correspond to more severe pathological states. Our goal was to identify a supervised latent factor that aligns with these pathological scores and uncover genes strongly associated with infarct-induced tissue damage. In each replicate, SuNSTAMP produced a supervised factor that showed a strong linear relationship with the SPaSE-derived pathological scores (Figure 4b). The Pearson correlation coefficient between the first factor of the *W* matrix (the supervised factor) and the pathological scores exceeded 0.80 in all replicates, highlighting the robustness and reproducibility of our framework across time points and biological replicates.

**Fig. 4.**
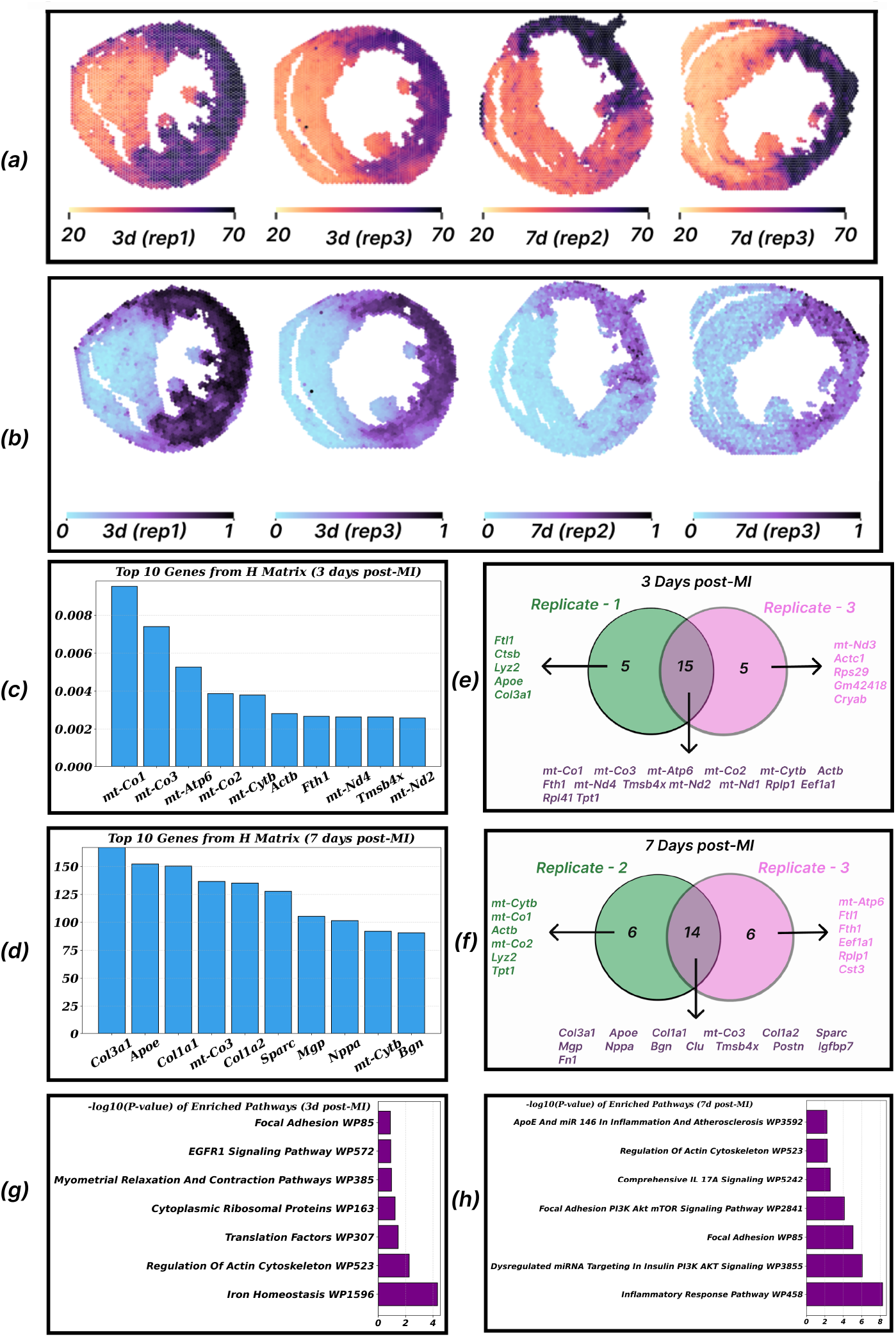
Application of SuNSTAMP to mouse Myocardial Infarction (MI) across two post-infarction time points and four biological replicates. *(a)* Spatial maps of pathological scores derived from the SPaSE pipeline across four replicates: Day 3 (Replicates 1 and 3) and Day 7 (Replicates 2 and 3). Higher scores indicate more severe infarct-induced tissue damage. *(b)* Scatter plots showing the strong linear relationship between SuNSTAMP’s supervised factor and the SPaSE-derived pathological scores in each replicate (Pearson *r >* 0.80 in all cases). *(c)* Top 10 genes contributing the most to the supervised factor found in 3 days post-MI (replicate-1), sorted in decreasing order of their values in the first row of the *H* matrix. *(d)* Top 10 genes contributing the most to the supervised factor found in 7 days post-MI (replicate - 2). *(e)* Venn diagram showing the overlap of top 20 genes identified by SuNSTAMP between the two Day-3 replicates (Replicates 1 and 3), with 15 genes shared. *(f)* Venn diagram for the two Day-7 replicates (Replicates 2 and 3), showing 14 overlapping genes among the top 20. *(g)* Statistical significance (-log10 P-value) of enriched pathways from the top 15 genes in 3 days post-MI replicate, with Iron Homeostasis WP1596 emerging as the most significant pathway. *(h)* Statistical significance (-log10 P-value) of enriched pathways from the top 15 genes in 7 days post-MI replicate. The Inflammatory Response Pathway WP458 emerges as the key pathway, in strong agreement with established findings in MI.

To validate the biological significance of genes prioritized by SuNSTAMP in the myocardial infarction (MI) dataset, we investigated the top 10 genes identified at both 3 days (Figure 4c) and 7 days (Figure 4d) post-infarction. Notably, the top gene sets show clear biological coherence with known pathological processes of MI, reflecting distinct temporal stages of disease progression.

At 3 days post-MI, the overlapping genes found amoung two replicates (replicate 1 and replicate 3) primarily consisted of mitochondrial-encoded components of the electron transport chain, including *mt-Co1, mt-Co2, mt-Co3, mt-Cytb, mt-Atp6, mt-Nd1, mt-Nd2*, and *mt-Nd4* (Figure 4e). These genes represent key subunits of respiratory complexes I, III, IV, and V, which are essential for oxidative phosphorylation. Dysregulation of these complexes has been directly linked to mitochondrial dysfunction, oxidative stress, and cardiomyocyte apoptosis during acute MI.

By Day 7, the overlapped genes (Figure 4f) identified by SuNSTAMP included key ECM (Extra-Cellular Matrix) remodeling and fibrosis-related genes—*Col1a1, Col3a1, Fn1, Postn, Sparc, Mgp, Nppa* — reflecting scar formation. This aligns with fibroblast trajectory studies in MI, where Day 7 fibroblasts exhibit myofibroblast-like phenotypes with increased ECM secretion and TGF-*β* signaling [23]. This findings support the validity of SuNSTAMP’s supervised factor in capturing biologically meaningful stage-specific gene activation in the context of MI.

Enrichment pathway analysis using Enrichr [18] further validated the biological meaningfulness and temporal specificity of SuNSTAMP-identified gene signatures. At 3 days post-MI, the top 15 genes were significantly enriched for Iron Homeostasis pathways (negative log p-value *>* 4) (Figure 4g), aligning with recent evidence that mitochondrial iron dysregulation is a critical early hallmark of myocardial ischemia-reperfusion injury [6]. This pathway enrichment corroborates our identification of mitochondrial electron transport chain genes, as excess mitochondrial iron accumulation during early MI increases reactive oxygen species production and exacerbates cardiomyocyte damage through ferroptosis and oxidative stress mechanisms.

By contrast, the 7-day gene signature showed pronounced enrichment for Inflammatory Response pathways (Figure 4h), reflecting the well-characterized temporal transition from acute metabolic dysfunction to sustained inflammatory tissue remodeling [13]. This temporal pathway progression—from iron dysregulation and mitochondrial dysfunction in acute injury to inflammatory-mediated tissue repair and fibrosis—precisely recapitulates the established pathophysiological timeline of post-infarction cardiac remodeling. The striking concordance between SuNSTAMP’s unsupervised gene prioritization and these functionally distinct, temporally-ordered biological processes demonstrates that our framework captures not merely statistical associations, but mechanistically relevant gene modules that drive disease progression.

This case study demonstrates the extendability of SuNSTAMP to tissues and diseases beyond neurodegeneration. Without altering the core algorithmic pipeline, SuNSTAMP successfully learned a disease-relevant latent factor that aligns closely with pathological annotations in myocardial infarction and enabled discovery of biologically meaningful gene signatures. These results underscore the broader applicability of our framework to diverse spatial transcriptomics datasets and disease types.

### Methodologies

#### Mathematical Formulation of Supervised NMF Update Rules

In the standard NMF, the matrix *X* is factorized into *W* (factor matrix) and *H*(coefficient matrix) matrices by minimizing the Frobenius norm of the difference between *X* and *WH*. Our supervised NMF framework modifies the objective function of standard NMF to include a supervision term that biases the first column or factor of *W* towards the supervision signal for making it a supervised factor. The modified objective function is:

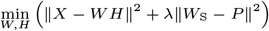

where

- *W*_S_ denotes the first factor of the *W* matrix.
- *P* denotes the supervision signal strength, computed using the spatial decay model of plaque signals in AD, and using SPaSE-based annotations in MI.

The multiplicative update rules for *W* and *H* were derived to iteratively minimize the objective function while ensuring non-negativity constraints.

To solve the optimization problem, we implemented an iterative algorithm that alternates between updating the *W* and *H* matrices. At each step, we ensure that all elements remain non-negative by applying multiplicative update rules. These update rules were carefully designed to accommodate the supervision term.

#### Derivation of Multiplicative Update Rules

The objective function is a combination of two terms. The first term denotes reconstruction error and the second term signifies the supervision term for a guided factorization process.

The loss function to optimize is:

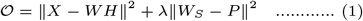

where

- *X* is an *n×m* gene expression matrix. Here n is the number of cells or spots and m is the number of genes in the gene expression matrix.
- *W* is an *n × k* matrix.
- *H* is a *k × m* matrix.
- k is the number of latent or hidden features in NMF.
- *λ* denotes the balance between reconstruction accuracy and adherence of supervised factor to supervision signal
- *W*_*S*_ is an *n × k* matrix where:

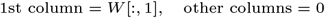
- *P* is the Supervision Matrix with dimension same as *W*_*S*_ where: 1st column = *P* [:, 1], Supervision values calculated for the cells,

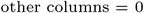

The objective function can be rewritten as:

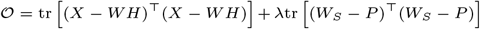

Simplifying further:

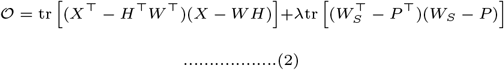

#### The role of Lagrange Multiplier

When deriving update rules for Non-negative Matrix Factorization (NMF) or similar problems, it’s essential to ensure that the factor matrices *W* and *H* remain non-negative (i.e., all elements *w*_*ik*_ ≥ 0 and *h*_*kj*_ ≥ 0). The Lagrange multiplier [27] method is a powerful technique used to incorporate these non-negativity constraints directly into the optimization problem.

Lagrange multipliers *ψ*_*ik*_ and *ϕ*_*kj*_ are introduced to incorporate the non-negativity constraints:

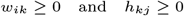

respectively. By introducing these multipliers, the problem is transformed into a Lagrangian, *ℒ*, which combines the original objective function with the constraints. The Lagrangian function *ℒ* is given by:

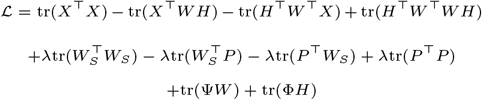

- Ψ and Φ: Matrices of Lagrange multipliers corresponding to the constraints on *W* and *H*.
- tr(Ψ*W*): Ensures that the non-negativity constraint *W* ≥ 0 is satisfied.
- tr(Φ*H*): Ensures that the non-negativity constraint *H* ≥ 0 is satisfied.

Taking the first order partial derivative of *ℒ* with respect to *W*,

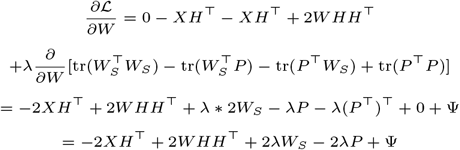

Setting 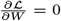, we get:

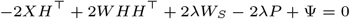

Taking the first order partial derivative of *ℒ* with respect to *H*:

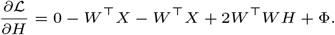

Setting 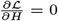,we get:

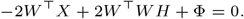

One of the key aspects of the KKT conditions [17] is the **complementary slackness condition**, which states that for each Lagrange multiplier *ψ*_*ik*_ and *ϕ*_*kj*_, either the constraint is active (i.e., *w*_*ik*_ = 0 or *h*_*kj*_ = 0), or the corresponding Lagrange multiplier is zero (i.e., *ψ*_*ik*_ = 0 or *ϕ*_*kj*_ = 0).

Using the KKT conditions, *ψ*_*ik*_ *ω*_*ik*_ = 0:

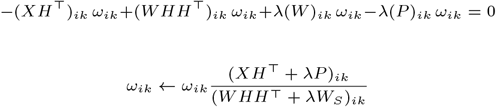

And,

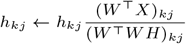

#### Final Results

Only the first column of *W* will get updated by the following rule:

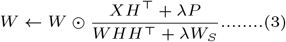

Other columns of *W* will get the update:

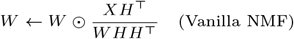

*H* will be updated by:

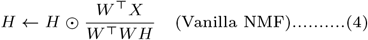

Theorem of Convergence

Regarding these two multiplicative update rules, we have the following theorem.

##### Theorem 1

The objective function *L* in (1) is non-increasing under the update rules in (3) and (4).

A detailed proof of the theorem can be found in the Supplementary Section 7. Our proof follows a similar approach to that presented in Lee and Seung’s [20] work on the original NMF. However, recent studies [5], [21] have shown that the multiplicative algorithm proposed by Lee and Seung [20] does not always guarantee convergence to a stationary point. In particular, Lin [21] introduced slight modifications to their algorithm, ensuring convergence. Since our update rules in (3) and (4) closely resemble those used in NMF, Lin’s modifications can be directly applied.

Moreover, when *λ* = 0, it is straightforward to verify that the update rules in (3) and (4) simplify to those of the original NMF formulation.

#### Hyperparameter Tuning via Cross Validation

We now focus on two critical hyperparameters. The first one is the minimal sufficient rank which maintains certain statistical significance, the number of latent features, and the second one is the best-fit *λ* value which modulates the extent of supervision applied during the multiplicative update steps. Here the parameter *λ* also balances between fitting the data and adhering to the supervision imposed by the first factor of the *W* matrix.

To get the best combination of rank and lambda, we employed a k-fold nested cross-validation strategy. K-fold nested cross-validation is a good approach that not only evaluates the model’s performance across multiple folds of the data but also ensures that the selection of hyperparameters is robust and generalizable [1].

In the outer loop of the nested cross-validation, the data was divided into k subsets, where the model was trained on k-1 subsets and validated on the remaining subset. This process was repeated k times, with each subset serving as the validation set once. The inner loop involved another k-fold cross-validation, where we performed a grid search over rank and lambda values to determine the combination that minimized the loss function.

Through this comprehensive approach, we identified the minimal rank and best-fit lambda values that best suited our model and maintained certain statistical significance.

#### Wilcoxon Rank-Sum Test for Minimal Sufficient Rank

After the k-fold nested cross-validation step, we have different rank-lambda combinations (particularly 21 different ranks starting from 5 to 25 and 26 different lambda values on different logarithmic scales) and their corresponding aggregated loss values. Now we proceed to analyze at which rank we can stop, as increasing the rank further shows diminishing improvements in the loss. To achieve this, we applied the Wilcoxon Rank Sum test [30] on consecutive pairs of ranks by comparing their point clouds, representing loss values for different lambda values.

The Wilcoxon rank-sum test is a non-parametric statistical test that does not assume any specific distribution for the data. It compares two independent samples to assess whether they come from the same distribution. In our context, the test compares the distributions of losses for consecutive ranks to determine whether there is a significant difference between them. If the difference is insignificant, it suggests that further increasing the rank may not provide meaningful improvements in the model’s performance.

We then plot the p-values from the Wilcoxon Rank Sum test against ranks. We focus on the peaks of the plot which imply increasing the rank further has no longer significant difference. We have adopted this method to get the minimal sufficient rank for our model.

#### Best-fit Lambda (*λ*)

After identifying the minimal sufficient rank using the Wilcoxon Rank-Sum test, we proceed to determine the best-fit value of the regularization parameter, *λ*. To achieve this, we evaluate each of the *λ* values considered during the nested k-fold cross-validation process by fixing the rank to the previously selected minimal sufficient rank. For each *λ* value, we retrain the model multiple times and compute the average loss to ensure stability and reduce the effect of randomness. In several cases, multiple *λ* values yielded similar average losses, making it difficult to distinguish a clear winner based solely on loss values.

To address this, we introduced an additional criterion based on the Pearson correlation coefficient between the supervision signal and supervised factor. Among the *λ* values with comparable losses, we selected the one which yielded a supervised factor that best captured the underlying signal complexity—i.e., the one with the highest Pearson correlation—indicating a better fit to capture the signal complexity.

## Discussion

The results from SuNSTAMP demonstrate its ability to extract biologically meaningful, spatially-informed gene expression patterns from high-resolution spatial transcriptomics data. In the Alzheimer’s Disease (AD) dataset, the supervised factor closely aligned with the spatial decay model of A*β* plaque signals, achieving a Pearson correlation of 0.92 (Figure 2c). This strong alignment underscores the effectiveness of focusing supervision on a single latent factor to capture disease-relevant spatial heterogeneity. Visual inspection of the spatial factor revealed clear anatomical patterns in cortical, hippocampal, and corpus callosum regions, which were not recapitulated by unsupervised NMF, highlighting the interpretability advantage of SuNSTAMP.

The top-ranked genes identified by our model—including *Ctsb, Cst3, Gfap*, and *Apoe*—exhibited progressive upregulation from 8- to 13-month-old AD mice, consistent with increasing plaque burden (Figure 3b). Cross-replicate analyses confirmed the stability and robustness of these gene rankings, with 14 genes consistently detected across independent samples and 70% pairwise ordering preserved (Supplementary Figure S3). Predictive validation further demonstrated that a compact set of 15-20 top genes suffices to achieve high AUC-ROC scores (≥ 0.90) in classifying spatial spots near versus far from plaques (Figure 3d). Orthogonal validation using the AlzMAP dataset reinforced these findings, showing elevated expression of the top genes in 12- and 18-month-old AD mice relative to control replicates [7].

Extending SuNSTAMP to the Myocardial Infarction (MI) dataset illustrated the generalizability of the framework beyond neurodegeneration. In this context, the supervised factor closely tracked SPaSE-derived pathological scores, with PCC values exceeding 0.80 across replicates and time points (Figure 4). The top genes discovered at 3 days post-infarction primarily represented mitochondrial components of the electron transport chain, consistent with acute oxidative stress and apoptosis, while Day 7 genes reflected extracellular matrix remodeling and fibrotic processes. These findings confirm that SuNSTAMP can identify stage-specific and biologically coherent gene programs across distinct tissues and disease models.

Overall, these results highlight the versatility, robustness and interpretability of SuNSTAMP. By leveraging spatially-informed supervision, the framework reliably identifies biologically meaningful latent factors and associated gene signatures, while maintaining reproducibility across replicates and disease contexts. The consistent detection of known pathological genes alongside novel candidates emphasizes its potential for uncovering underexplored mechanisms in complex tissues.

SuNSTAMP provides a generalizable computational approach for spatially guided, supervised factorization of multimodal transcriptomic data. In AD, it effectively captures plaque-associated gene expression patterns and predicts spatial proximity to A*β* plaques. In MI, it uncovers temporally and spatially coherent gene programs related to infarct progression. These results demonstrate that SuNSTAMP is a broadly applicable tool for integrating disease-relevant supervision into spatial transcriptomics analyses, facilitating the discovery of robust and interpretable gene signatures across diverse biological contexts.

## Acknowledgements

The supervised Non-negative Matrix Factorization (NMF) model used in this study was implemented based on the NMF module from the scikit-learn library (https://github.com/scikit-learn/scikit-learn/blob/main/sklearn/decomposition/_nmf.py). Faria Binta Awal is supported by Undergraduate Research Grant from RISE, BUET. M Saifur Rahman is partially supported by Basic Research Grant from BUET.

## Data and Code Availability

The datasets utilized in this study include both Alzheimer’s Disease (AD) and Myocardial Infarction (MI) models.

For AD, STARmap PLUS sequencing data are available on the Single-Cell Portal at https://singlecell.broadinstitute.org/single_cell/study/SCP1375 and on Zenodo at https://doi.org/10.5281/zenodo.7332091. Further details on the dataset can be found in the original publication https://www.nature.com/articles/s41593-022-01251-x.

For MI, the sc/snRNA-seq data and spatial transcriptomic sequencing data have been deposited in the Gene Expression Omnibus under accession number GSE214611 (https://www.ncbi.nlm.nih.gov/geo/query/acc.cgi?acc=GSE214611). All other data supporting the findings of that study are included in the original article [4] and its associated files.

Source code for SuNSTAMP is publicly available on GitHub (https://github.com/f12-mou/SuNSTAMP). This repository contains all scripts necessary to replicate the findings of this study.

## Supplementary

**Table S1.**
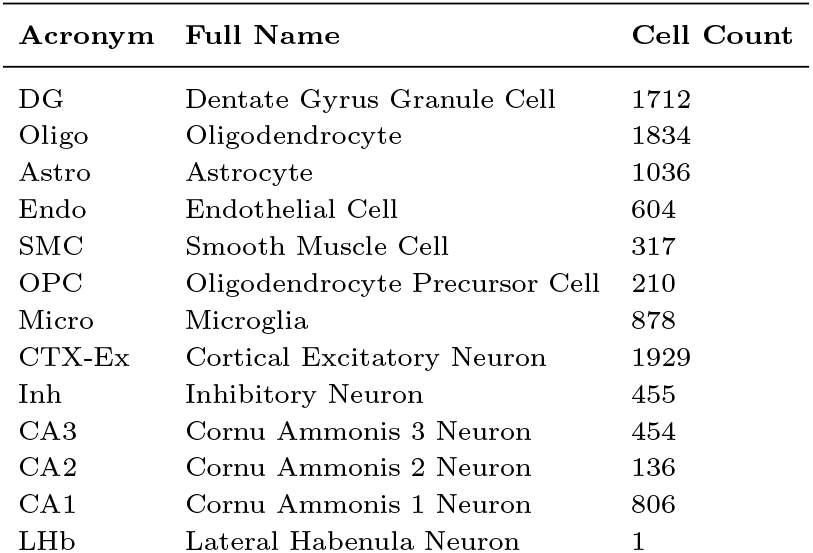
Cell type acronyms, full names, and their counts across all brain regions in AD 13 months old mouse model (replicate-1).

### Proof of Theorem 1

#### Loss Function

Our objective function was

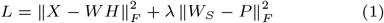

where,

- *X ∈* ℝ^*n×m*^ is the gene expression matrix,
- *W ∈* ℝ^*n×k*^ is the factor matrix,
- *H ∈* ℝ^*k×m*^ is the coefficient matrix,
- *W*_*S*_ *∈* ℝ^*n×k*^ is the matrix where
  - *W*_*S*_’s 1st column = *W* ‘s 1st column,
  - *W*_*S*_’s other columns = 0,
- *P* is the supervision signal matrix.

The objective function of SuNSTAMP in (1) can not have negative values. So, our objective function is bounded from below by zero.

The update rules are:

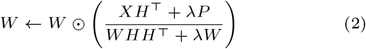

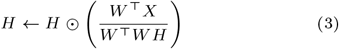

#### Proof

Proof: To prove Theorem 1, we need to show that *L* is nonincreasing under the update rules of (2) and (3). As the 2nd term of the loss function *L* is only related to *W*, we have the same update formula for *H* (eqn (3)) as in the original NMF. Thus, we can use the convergence proof of NMF to show that *L* is non-increasing under the update rule for *H*. We only need to prove that *L* is non-increasing under the update step of *W* according to equation (2).. We will follow a procedure similar to that described in [19]. Our proof will make use of an auxiliary function similar to that used in the expectation maximization algorithm [11].

#### Definition of Auxiliary Function

*G*(*w, w*^*′*^) is an auxiliary function for *F* (*w*) if the following conditions are satisfied:

- *G*(*w, w*^*′*^) ≥ *F* (*w*)
- *G*(*w, w*) = *F* (*w*)

An auxiliary function *G*(*w, w*^*′*^) is a function that helps in optimizing another function *F* (*w*) by providing an easier way to iteratively minimize *F* (*w*). The conditions described here showed that *G*(*w, w*^*′*^) serves as an upper bound to *F* (*w*), but at *w* = *w*^*′*^, they are equal.

The auxiliary function is also very useful for the following lemma:

##### Lemma - 1

If *G* is an auxiliary function of *F*, then *F* is non-increasing under the update:

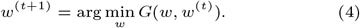

Proof of the Lemma - 1

From the definition of an auxiliary function *G*(*w, w*^*′*^), we know that:

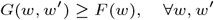

This means *G* always upper-bounds *F*.

Applying this at *w* = *w*^(*t*+1)^, we get:

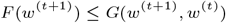

This follows because *G* is an upper bound on *F*. By definition, we chose *w*^(*t*+1)^ to minimize *G*(*w, w*^(*t*)^):

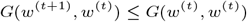

This means that replacing *w*^(*t*+1)^ with *w*^(*t*)^ would have given a larger or equal value.

Since the auxiliary function satisfies *G*(*w, w*) = *F* (*w*), we substitute:

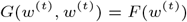

This is just the definition of an auxiliary function.

Combining these results:

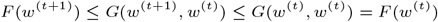

This shows that *F* (*w*) is nonincreasing with each update step.

Now, we have to show that the update step for *W* in equation (2) is exactly the update in (4) with a proper auxiliary function.

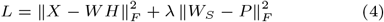

Considering any element *w*_*ab*_ in *W*, we use *F*_*ab*_ to denote the part of *L* which is only relevant to *w*_*ab*_.

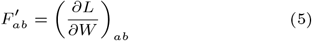

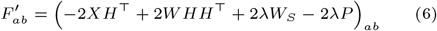

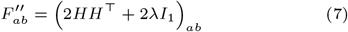

Here, *I*_1_ is the identity matrix, which we are considering to be *n × k* with 1 at the 1st row, 1st column, and other values are 0.

#### Design of the Auxiliary Function

The auxiliary function 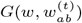 is defined as:

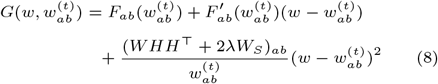

##### Lemma - 2

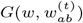 is an auxiliary function for *F*_*ab*_, the part which is only relevant to *W*_*ab*_.

Proof of Lemma - 2

Since *G*(*w, w*) = *F*_*ab*_(*w*) is obvious, we only need to show that

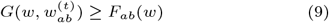

We compare the Taylor series expansion [12] of *F*_*ab*_(*W*):

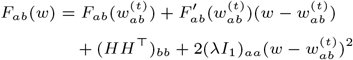

Here,

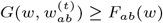

is equivalent to

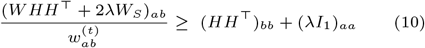

We will show the proof of the inequality in two parts:

- 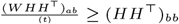
- 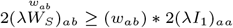

1st part,

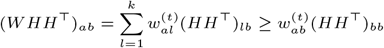

As, the LHS is a sum of some terms and RHS is one of those terms, therefore, LHS ≥ RHS

For the second part,

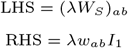

if *b* = 1, then

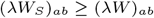

otherwise the values become 0.

So, it holds 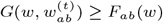.

Therefore, *G* is an auxiliary function of *F*.

*G* was minimized for a given *w*^(*t*)^ to get the next *w*^(*t*+1)^. So, *F* is non-increasing.

**Fig. S1.**
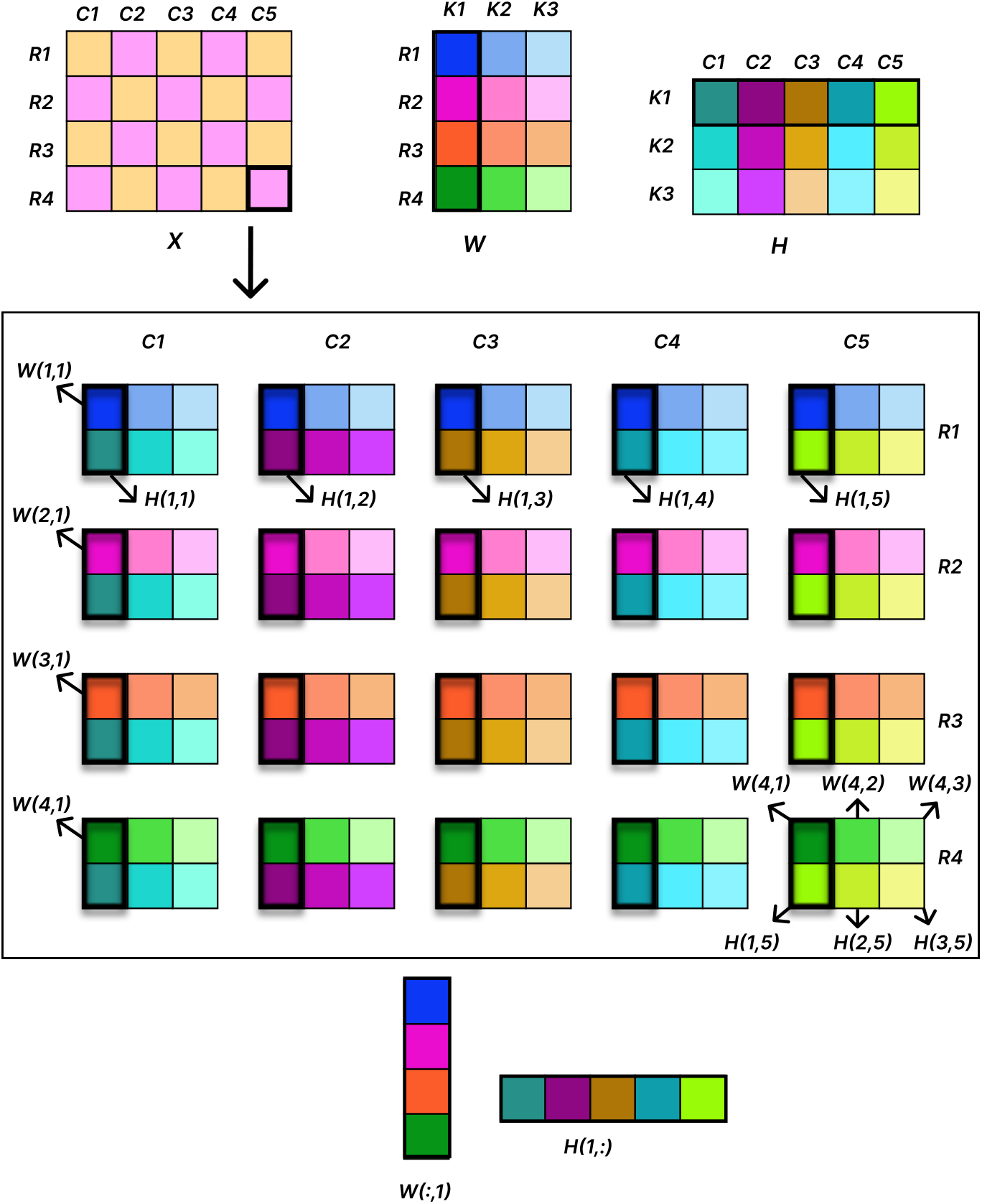
Illustration of the non-negative matrix factorization (NMF) of a data matrix *X* into the product of two lower-dimensional non-negative matrices: *W* and *H*. Each cell of these matrices is visually differentiated using distinct colors to denote individual numerical values. The original matrix *X*, shown on the left, consists of 4 samples (rows) and 5 features or genes (columns). The decomposition into three latent factors (rank, k=3) introduces an interpretable intermediate structure. The matrix *W* captures how much each sample is associated with each factor. The matrix *H* describes how strongly each factor activates each feature (gene). The reconstruction of *X* via multiplication of *W* and *H* is demonstrated. For example, for the marked cell in *X*(4, 5), the reconstruction involves *W* (4, 1) * *H*(1, 5) + *W* (4, 2) * *H*(2, 5) + *W* (4, 3) * *H*(3, 5). This decomposition is visualized as three color-coded elementwise products. It is visually clear how a single cell in *X* is explained by three (factor, coefficient) pairs. We then select one such (factor, coefficient) pair per cell to apply supervision on the factor, as demonstrated in the picture, the first pair of each cell. Thus, we want the first factor to allign with a specific supervision signal, and thereby get the distribution of coefficients of features (or genes) towards this signal. Through this selection process across all cells, a simple and quite obvious pattern emerges: supervision applied to a specific factor (say, factor *j*) naturally focuses our attention on the *j* − *th* row of matrix *H*.

**Fig. S2.**
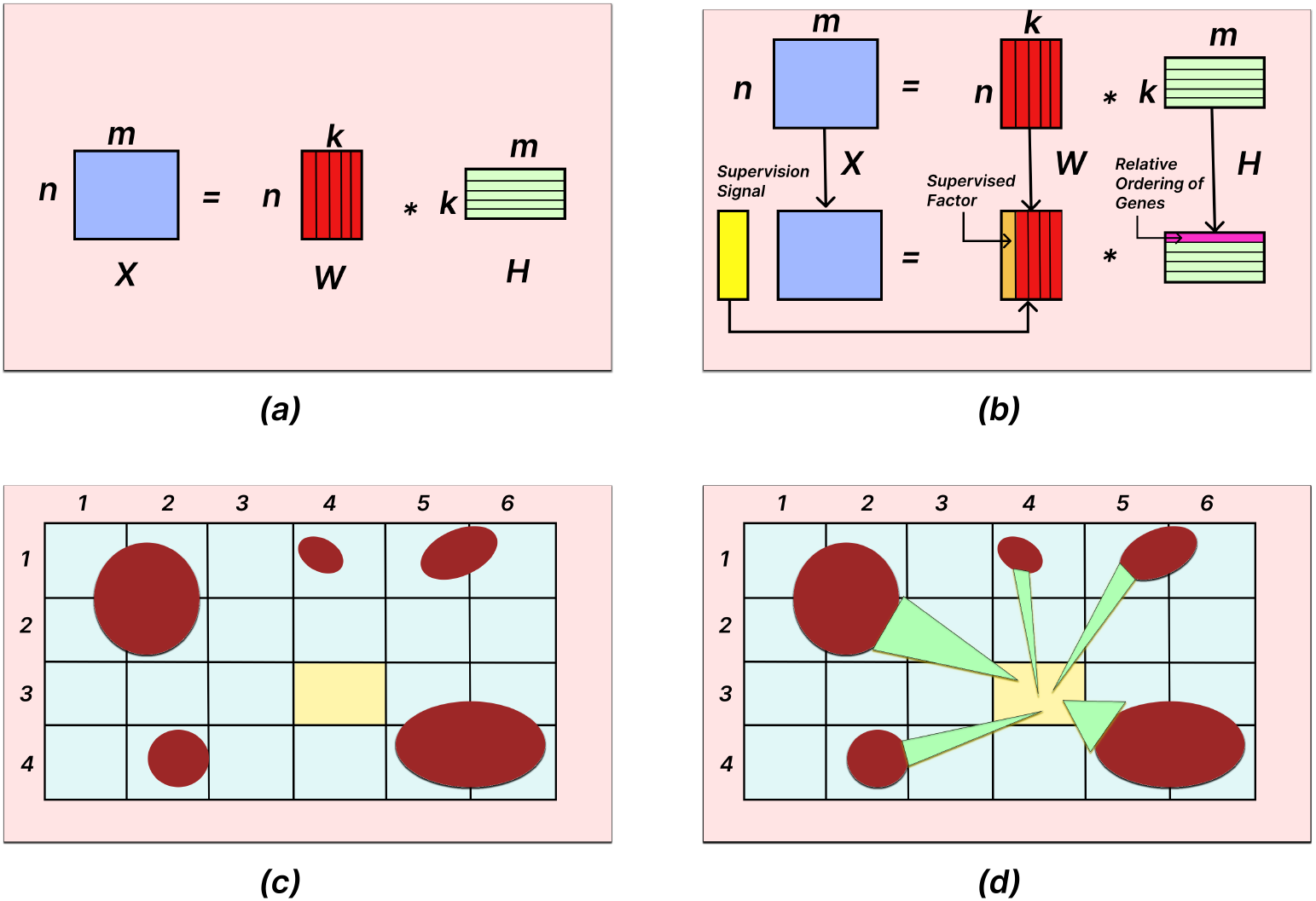
*(a)* Illustration of Non-negative Matrix Factorization (NMF). The input data matrix *X*_*n*\**m*_ (2D matrix that has n observations and m features) is decomposed into two non-negative matrices: *W*_*n*\**k*_ (basis matrix capturing latent factors) and *H*_*k*\**m*_ (coefficient matrix representing the contribution of each feature to the factors). Here *k* denotes rank of NMF which is the number of latent factors extracted from the data matrix, *X*_*n*\**m*_. *(b)* We restrict the supervision signal (represented by yellow column vector) to *a single factor* keeping all the other factors unaffected. This enables better signal-specific interpretability of the corresponding row in the coefficient matrix (*H*). *(c)* In this grid-based representation of tissue, red ellipses denote A*β* plaques, and each grid cell represents a spatial unit (e.g., a spot or cell). The yellow-highlighted cell at coordinate (3,4) contains no plaque within its boundary and is thus assigned a plaque intensity of zero in a spot-centric model. However, it is surrounded by large neighboring plaques, suggesting potential biological influence on it. This rigid boundary-based assumption fails to capture the continuous spatial nature of plaque effects. *(d)* Each plaque exerts an influence that gradually decays with distance between the plaque itself and a cell, allowing nearby cells—such as the yellow-marked cell at (3,4) to inherit partial intensity based on proximity. This distance-aware modeling better reflects the gradient-like diffusion of A*β* plaque effects (green triangles whose base denotes higher plaque intensity and peak denotes lower intensity) and improves the detection of plaque-associated gene expression beyond rigid spot boundaries.

**Fig. S3.**
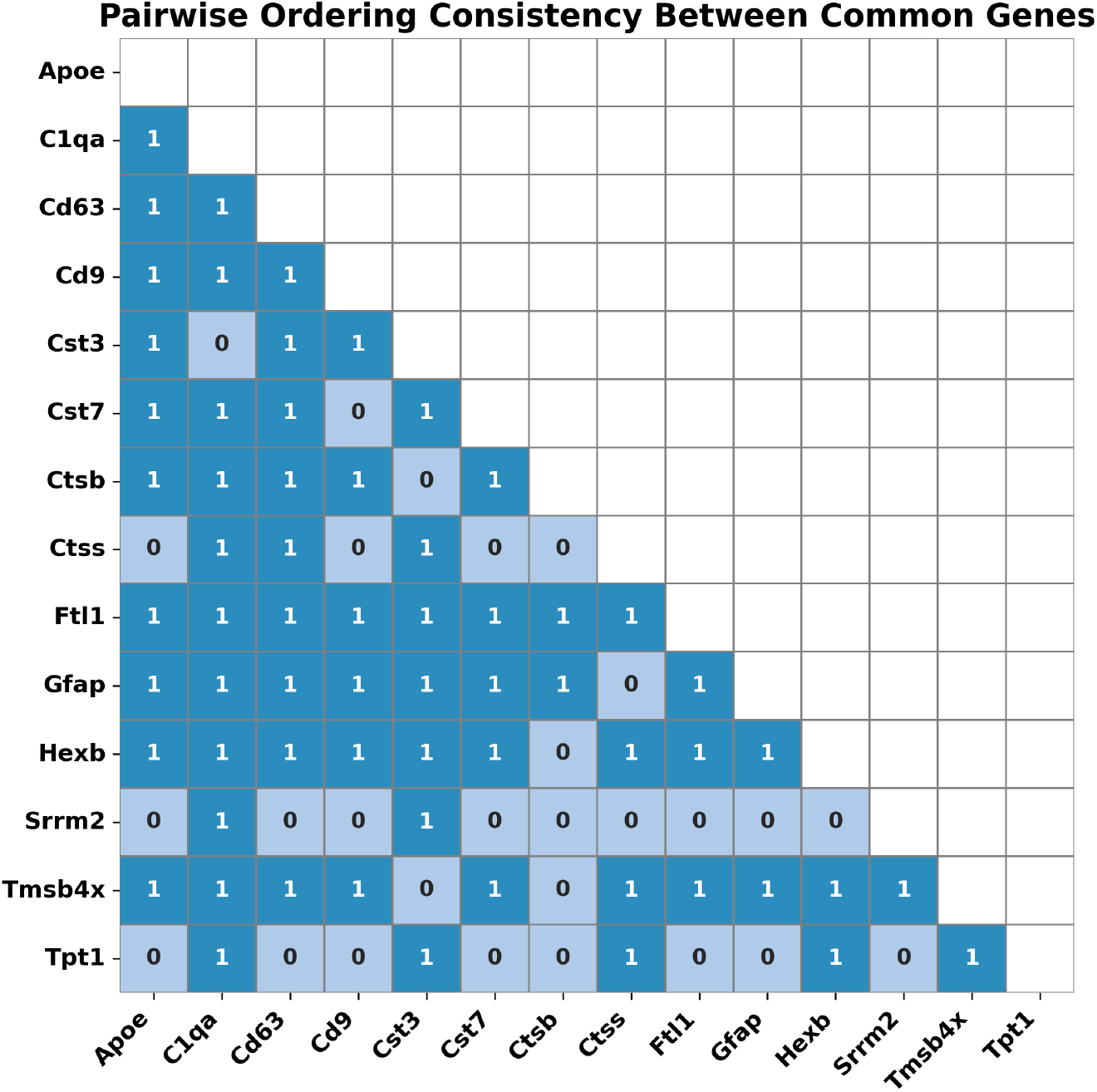
Pairwise ordering consistency matrix for the 14 genes commonly identified in both 13-month-old replicates. Each cell in the lower triangle compares the relative ranking of a gene pair across replicates. A value of 1 (blue) indicates consistent ordering—that is, the higher-ranked gene in one replicate also ranks higher in the other. A value of 0 (light blue) indicates inconsistent ordering. This matrix reveals that 70% of all gene pairs maintain consistent ordering.

**Fig. S4.**
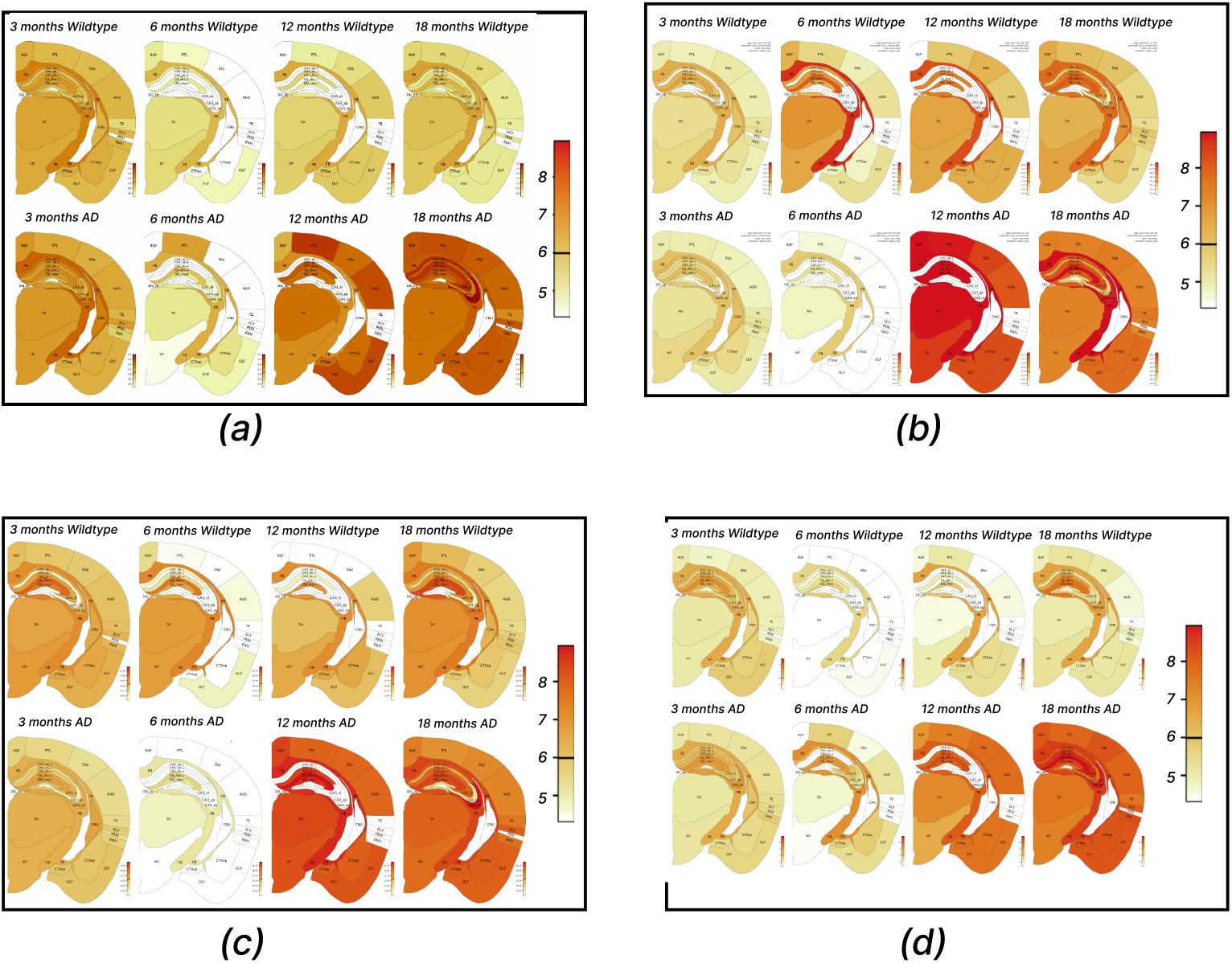
Expression profiles of selected top-ranked genes *(a)Cst3, (b)Ctsb, (c)Apoe, (d) Gfap* across control and Alzheimer’s disease (AD) mouse brains at 3, 6, 12, and 18 months from an independent dataset [7]. Each gene’s expression is visualized with control (top row) and AD (bottom row) across these 4 different time points. Warmer color (red) indicate higher expression levels. Notably, elevated expression in AD mice at 12 and 18 months supports the relevance of SuNSTAMP-identified genes in later stages of disease progression.

## Notes

https://github.com/f12-mou/SuNSTAMP

### Competing Interest Statement

The authors have declared no competing interest.

